# *In vivo* studies of glucagon secretion by human islets transplanted in mice

**DOI:** 10.1101/2019.12.15.876920

**Authors:** Krissie Tellez, Yan Hang, Xueying Gu, Roland W. Stein, Seung K. Kim

**Author notes:** These authors contributed equally (Krissie Tellez, Yan Hang). Communicating author: Seung K. Kim, 279 Campus Drive, Stanford, CA 94305.

## Abstract

Relatively little is known about regulated glucagon secretion by human islet α cells compared to insulin secretion from β cells, despite conclusive evidence of dysfunction in both cell types in diabetes mellitus. Distinct insulin sequences in humans and mice permit *in vivo* studies of β cell regulation after human islet transplantation in immunocompromised mice, whereas identical glucagon sequences prevent analogous *in vivo* measures of glucagon output from human α cells. We used CRISPR/Cas9 genome editing to remove glucagon-encoding codons 2-29 in immunocompromised (*NSG*) mice, preserving production of other proglucagon-derived hormones, like Glucagon-like-peptide 1. These *NSG-*Glucagon knockout (*NSG-GKO*) mice had phenotypes associated with glucagon signaling deficits, including hypoglycemia, hyperaminoacidemia, hypoinsulinemia, and islet α cell hyperplasia. *NSG-GKO* host metabolic and islet phenotypes reverted after human islet transplantation, and human islets retained regulated glucagon and insulin secretion. *NSG-GKO* mice provide an unprecedented resource to investigate unique, species-specific human α cell regulation *in vivo*.

---

Pancreatic islet α and β cells play an important role in maintaining euglycemia by secreting peptide hormones in response to glucose and other blood metabolites. In healthy β cells, hyperglycemia triggers insulin secretion, which promotes glucose uptake and glycogenesis or adipogenesis in ‘insulin-target’ organs. In contrast, α cell glucagon secretion, stimulated by hypoglycemia, amino acids, and autonomic nerve inputs, leads to glucose mobilization by promoting glycogenolysis and gluconeogenesis in ‘glucagon-target’ organs, like liver^1^. Impaired regulation or output of insulin and glucagon by human β cells and α cells underlies development and progression of diabetes mellitus. Thus, intensive efforts are focused on determining the physiological and pathological mechanisms governing human islet α cell and β cell function.

Recent studies reveal that human and mouse islet cells have differences in cellular composition, molecular regulation, physiological control, intra-islet cell interactions and other crucial properties^2-5^, motivating increased research focus and resource generation in human islet biology. Transplantation of human islets in immunocompromised mice, like the *NOD.Cg-Prkdc*^*scid*^*Il2rg*^*tm1Wjl*^*Sz* mice (*NSG*) strain^6,7^, has emerged as an important strategy for assessing human islet β cell function *in vivo*^8-10^. Unlike distinct human and mouse insulins, the mature glucagon sequence in these species is identical, precluding accurate quantification of circulating human islet-derived glucagon secretion in mice and limiting studies of human α cells in transplantation-based models. Thus, development of immunocompromised mouse strains that permit detection of human glucagon in mice and *in vivo* studies of transplanted human islet α cell function could be transformative by enabling mechanistic analysis under normal and pathophysiological conditions.

Genetic targeting to eliminate endogenous glucagon production in mice could permit *in vivo* quantification of glucagon output by transplanted human islets. The *Gcg* gene encodes proglucagon, a prohormone expressed and differentially processed in islet α cells, gut enteroendocrine cells, and the central nervous system to produce multiple distinct peptide hormone products, including glucagon, oxyntomodulin, glicentin, glucagon-like peptide-1 (GLP-1) and glucagon-like peptide 2 (GLP-2)^11^. Differential proglucagon processing depends on co-expression of the prohormone convertase (PC) enzymes. In pancreatic islet α cells, PC2 enables cleavage of proglucagon into the mature 29 amino acid glucagon protein, which is entirely encoded by *Gcg* exon 3^12^. PC1/3 expression in enteroendocrine cells permits cleavage of proglucagon into other products, including GLP-1, a secreted incretin hormone that enhances postprandial insulin output by islet β cells^13-17^.

Glucagon production in mice has been eliminated by targeted mutation of the *Gcg* gene to generate the *Gcg*^gfp^ allele; adult homozygous *Gcg*^gfp/gfp^ mice appeared normoglycemic, but exhibited α cell hyperplasia and hypoinsulinemia^18^. In contrast, loss of glucagon signaling in mice with a glucagon receptor (*Gcgr*) deletion, by Gcgr antibody inactivation^19-22^, or elimination of PC2^23, 24^ results in more extensive metabolic phenotypes reflecting impaired gluconeogenesis and glycogenolysis, including elevated circulating amino acid levels and basal hypoglycemia. Therefore, the milder metabolic phenotypes of *Gcg*^gfp/gfp^ mice^18^ likely reflect the combined loss of mature glucagon and other products of proglucagon. Here we generated immunocompromised mice that solely lack mature glucagon coding sequences (*NSG-GKO*) and demonstrate the utility of *NSG-GKO* mice for analyzing human islet α cell function *in vivo*.

## Results

### Generation of NSG-GKO mice

To develop mice that permit transplantation of human islets and detection of human glucagon, we used CRISPR/Cas9 genome editing in NSG-derived oocytes to create an in-frame deletion of nucleotides from *Gcg* exon 3, which encode mature glucagon (**Figures 1A and S1A**). This strategy should preserve production of metabolic regulators derived from the proglucagon carboxy-terminus, including GLP-1. After generating candidate founder mice, genotype screening identified one *NSG* founder harboring an in-frame 93 base pair (bp) deletion of the 3’ end of *glucagon* exon 3. This in-frame deletion includes the elimination of 84 nucleotides encoding amino acids 2-29 of mature glucagon (**Figures 1A-B and S1A)**. Subsequent breeding of heterozygous F1 mice produced viable, fertile homozygous *NSG-GKO* mice that were born at a rate of 22.1% (compared to 32.8% wild type, and 45.1% heterozygous; n= 122 mice). While three-week-old male and female *NSG-GKO* mice weighed significantly less than *NSG* control littermates, no difference in body weight was detected in adult mice at eight weeks of age (**Figure S1C**). This transient reduction of body mass in *NSG-KO* mice likely reflects a combination of reduced circulating insulin levels in *NSG-GKO* mice (see below), and possibly unrecognized roles for glucagon in development^25-28^. Thus, beginning at eight weeks of age, *NSG-GKO* mice of both sexes were characterized for phenotypes associated with glucagon signaling loss.

**Figure 1.**
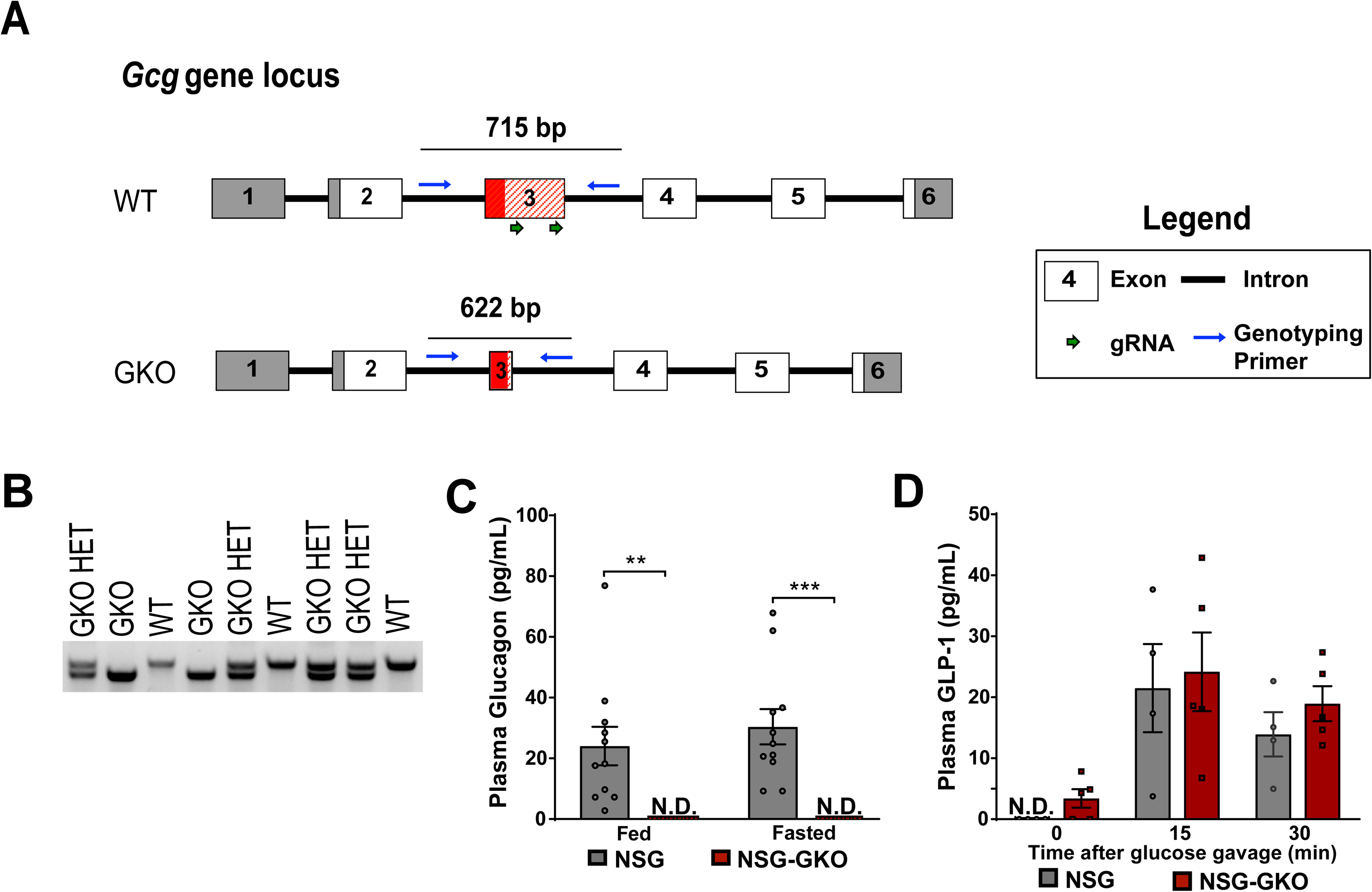
Generation of *NSG-GKO* mice. (**A**) Schematic showing *Glucagon* (*Gcg*) gene structure, guide RNA (gRNA) targeting sites (green arrows), and genotyping primers (blue arrows). Exon 3 is highlighted in red with the portion encoding mature glucagon marked by hatch lines. (**B**) Representative genotyping PCR of *NSG-GKO* mice following a heterozygous *NSG-GKO* cross. (**C**) Plasma glucagon levels in 2-3 month old *NSG-GKO* and *NSG* mice during *ad libitum* feeding (*P*= 0.0007 by two-tailed Student’s t-test) or after a 3-hour fast (*P*= 0.0001 by two-tailed Student’s t-test) (*NSG* mice, n= 11; *NSG-GKO* mice, n= 12). (**D**) Plasma GLP-1 levels from 2.5-3 month old *NSG-GKO* and *NSG* controls following oral glucose challenge (*NSG* mice, n= 4; *NSG-GKO* mice, n= 5). Data are represented as mean of biological replicates ± SEM. * *P* ≤ 0.05, ** *P* ≤ 0.01, *** *P* ≤ 0.001. N.D.= not detected. See also Figure S1.

To validate elimination of mature glucagon, plasma glucagon levels were measured from *ad libitum* fed and fasted *NSG-GKO* and *NSG* control mice. Unlike control *NSG* mice, plasma glucagon was undetectable in *NSG-GKO* mice in both fed and fasted states (**Figure 1C**). By contrast, plasma GLP-1 levels and excursion following an oral glucose tolerance test were similar between *NSG-GKO* and *NSG* control littermates, despite the increased glucose tolerance seen in NSG-GKO mice (**Figures 1D and S1D**). Consistent with this, antibodies that detect mature glucagon (GCG1-29) did not label islet α cells in *NSG-GKO* mice, whereas these cells were readily identified with an antibody that detects proglucagon (**Figure S1B**). Thus, our CRISPR-based strategy successfully eliminated glucagon production while preserving GLP-1 output in *NSG-GKO* mice.

### Transplanted human islets retain regulated glucagon secretion in NSG-GKO mice

To assess the possibility of measuring circulating glucagon from human islets in *NSG-GKO* mice, we transplanted human islets from previously-healthy donors under the renal capsule of *NSG-GKO* mice (*NSG-GKO Tx* mice) (**Figure 2A**). Plasma glucagon was detectable in *NSG-GKO Tx* mice two weeks after transplantation and thereafter for at least fourteen weeks, at which point *NSG-GKO Tx* mice were sacrificed for tissue analysis. Four weeks after human islet transplantation, plasma glucagon levels in fasted *NSG-GKO Tx* and control *NSG* mice were remarkably similar, but circulating glucagon remained undetectable in sham-transplanted *NSG-GKO* mice (**Figure 2B**).

**Figure 2.**
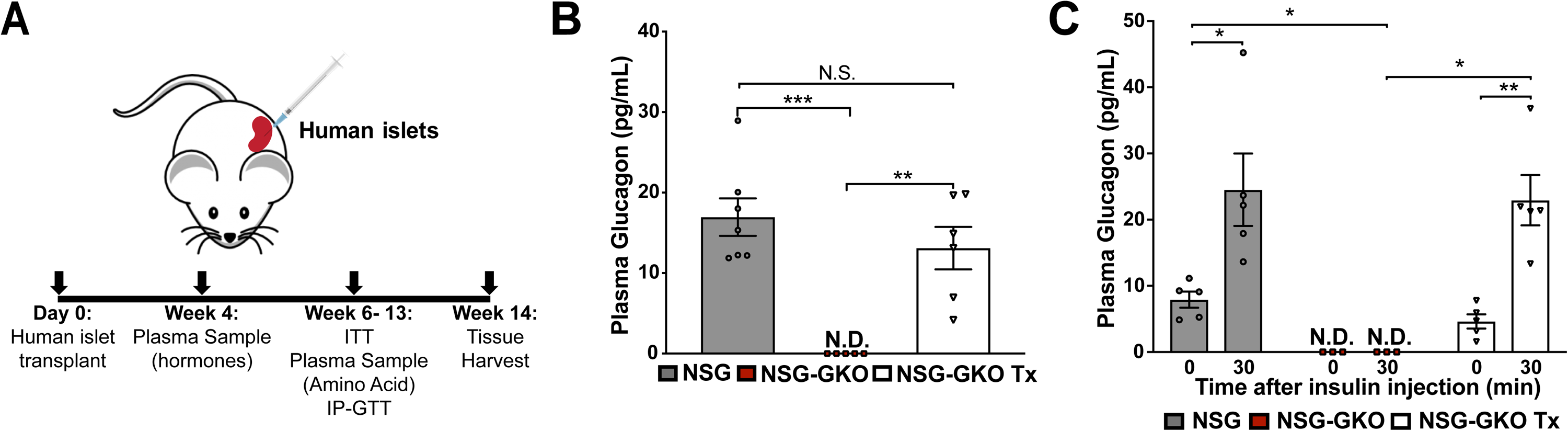
Transplanted human islets retain regulated glucagon secretion in *NSG-GKO* mice. Human islets were transplanted under the renal capsule of *NSG-GKO* mice. Mice were then examined for presence of glucagon in the circulation and regulation of glucagon secretion from transplanted islets. Data are from *NSG* control, *NSG-GKO*, and *NSG-GKO* mice post-transplantation (*NSG-GKO* Tx). (**A**) Schematic of islet transplantation and phenotyping schedule. (**B**) Fasted plasma glucagon levels (*NSG* vs. *NSG-GKO P*= 0.0002 and *NSG-GKO* vs. *NSG-GKO* Tx *P*= 0.0031 by one-way ANOVA, with Tukey’s multiple comparisons test) (*NSG* mice, n= 7; *NSG-GKO* mice, n= 5; *NSG-GKO* Tx mice, n= 6). (**C**) Mice were challenged with human insulin; glucagon response to acute hypoglycemia was measured from plasma at 0 and 30-minutes post-insulin injection (0’ vs 30’: *NSG P*= 0.0199 and *NSG-GKO* Tx *P*= 0.0091 by two-tailed Student’s t-test. 30’: *NSG* vs. *NSG-GKO P*= 0.0130 and *NSG-GKO* v. *NSG-GKO* Tx *P*= 0.0189 by one-way ANOVA, with Tukey’s multiple comparisons test) (*NSG* mice, n= 5; *NSG-GKO* mice, n= 3; *NSG-GKO* Tx mice, n= 5). Data are represented as mean of biological replicates ± SEM. N.D.= not detected. * *P* ≤ 0.05, ** *P* ≤ 0.01, *** *P* ≤ 0.001. See also Figure S2.

To assess dynamic regulation of glucagon secretion from human α cells in *NSG-GKO Tx* mice, we measured glucagon secretion *in vivo* after an intraperitoneal insulin challenge, which elicits transient hypoglycemia. Reduced blood glucose levels stimulate α cell glucagon secretion^1, 29^, and, as expected, acute hypoglycemia was accompanied by increased circulating glucagon levels in *NSG* controls (**Figures 2C and S2**). By contrast, insulin challenge and hypoglycemia elicited no glucagon output in sham-transplanted *NSG-GKO* mice (**Figures 2C and S2**). Like in *NSG* controls, circulating human islet-derived glucagon levels increased upon induction of hypoglycemia in *NSG-GKO Tx* mice (**Figures 2C and S2**). As transplanted human islets are the sole source of circulating glucagon in *NSG-GKO Tx* mice, we conclude that insulin challenge and ensuing transient hypoglycemia evoked glucagon secretion by human α cells in these mice, an *in vivo* response not previously reported for human islets transplanted in mice. These results suggest human α cell mechanisms governing regulated glucagon secretion remained intact after transplantation.

### Human islets establish a glucagon-signaling axis that corrects liver phenotypes in NSG-GKO mice

Glucagon signaling is an essential regulator of hepatic amino acid metabolism, gluconeogenesis, and glycogenolysis^20, 30, 31^. Consistent with prior mouse models of impaired glucagon signaling from glucagon or glucagon receptor deficiency^20-22^, we observed increased total plasma amino acid levels in *NSG-GKO* mice compared to *NSG* control littermates (**Figure 3A**). Moreover, measures of individual amino acids revealed specific amino acids that showed a significant increase in *NSG-GKO mice*, including arginine, alanine, threonine, serine, leucine, and glycine (**Figures 3B and S3**). To examine the impact of glucagon loss on gluconeogenesis and amino acid metabolism, hepatic gene expression of enzymes involved in gluconeogenesis (*G6pc and Pepck)* and amino acid metabolism (*Tat, Oat, Nnmt*, and *Gls2*) were measured by qPCR. Hepatic expression of *G6pc, Tat, Oat, Nnmt*, and *Gls2* was significantly decreased in *NSG-GKO* mice compared to *NSG* controls (**Figure 3D**). To assess differences in glycogenolysis, liver glycogen levels were measured from fasted NSG-GKO and NSG control mice. We observed a trend of increased average hepatic glycogen levels in NSG-GKO mice compared to NSG control (**Figure 3C**). Together, these data suggest that, like in previous studies of glucagon signalling loss^20-22, 24, 38^, gluconeogenesis and amino acid metabolism are impaired in *NSG-GKO* mice.

**Figure 3.**
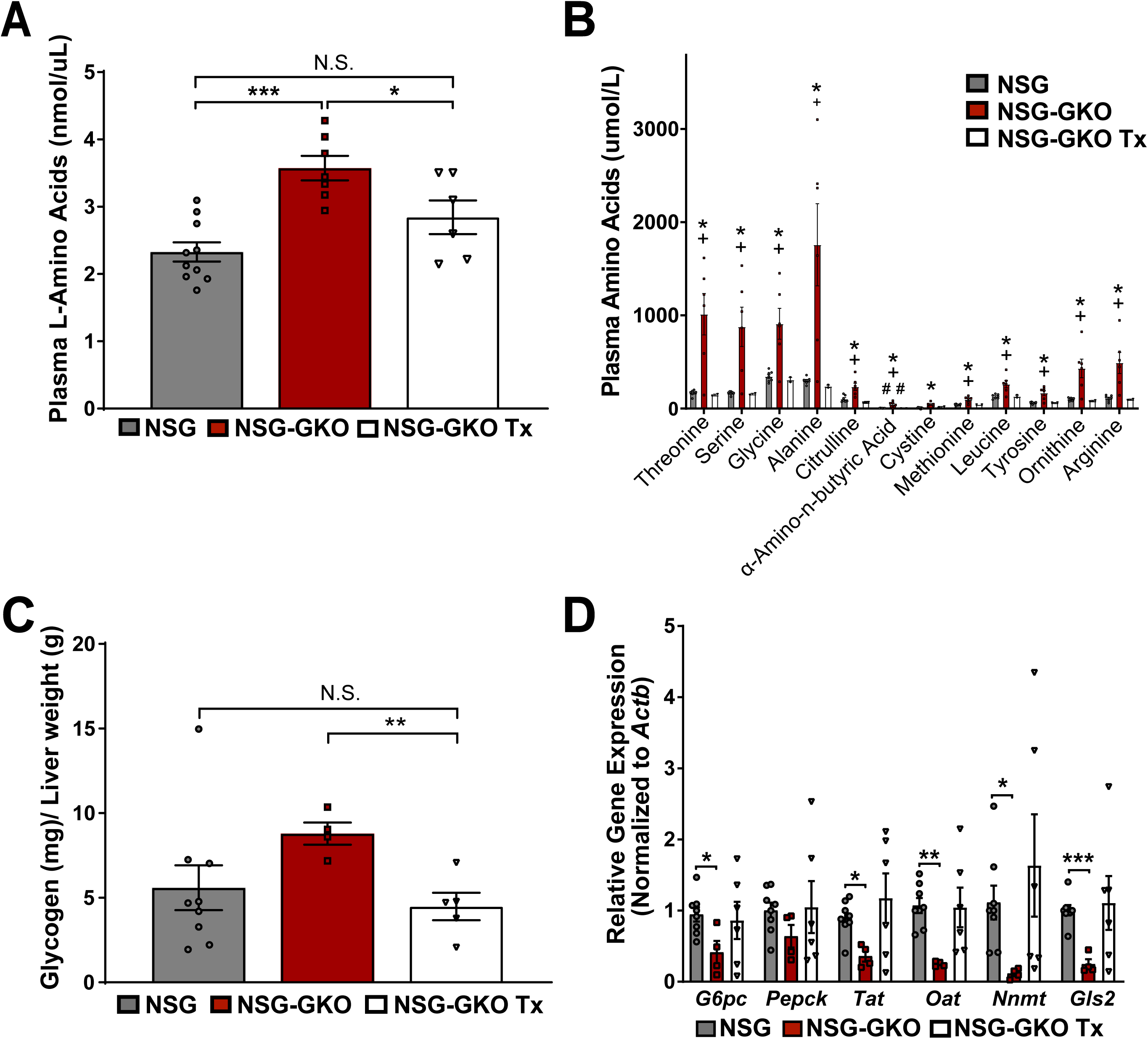
Human islet transplantation establishes a glucagon-signaling axis that corrects liver phenotypes in *NSG-GKO* mice. Mice were examined for liver phenotypes associated with glucagon loss following transplantation of human islets. Data are from *NSG* control, *NSG-GKO*, and *NSG-GKO* mice post-transplantation (*NSG-GKO* Tx). (**A**) Total plasma amino acids (*NSG* vs. *NSG-GKO P*= 0.0002 and *NSG-GKO* vs. *NSG-GKO* Tx *P*= 0.0442 by one-way ANOVA, with Tukey’s multiple comparisons test ANOVA) (*NSG* mice, n= 10; *NSG-GKO* mice, n= 7; *NSG-GKO* Tx mice, n= 6). (**B**) Concentration of individual plasma amino acids that showed significant change in NSG-GKO mice by two-way ANOVA, with Tukey’s multiple comparisons test (*P* values listed in Table S2) (*NSG* mice, n= 7; *NSG-GKO* mice, n= 6; *NSG-GKO* Tx mice, n= 2). (**C**) Liver glycogen quantification from the left lobe (*P*= 0.0054 by two-tailed Student’s t-test) (*NSG* mice, n= 9; *NSG-GKO* mice, n= 4; *NSG-GKO* Tx mice, n= 5). (**D**) Gene expression in the left liver lobe of indicated genes (significant *P* values generated by two-tailed Student’s t-test, with Holm-Sidak’s multiple comparisons test are listed in Table S2) (*NSG* mice, n= 8; *NSG-GKO* mice, n= 4; *NSG-GKO* Tx mice, n= 6). Data are represented as mean of biological replicates ± SEM. * *NSG* vs. *NSG-GKO* mice, ^+^ *NSG-GKO* vs. *NSG-GKO Tx* mice, and ^#^ *NSG* vs. *NSG-GKO* Tx mice (**B** and **D**). ^#, +^, or * *P* ≤ 0.05, ^##, ++^, or **, *P* ≤ 0.01, ^###, +++^ or ***, *P* ≤ 0.001. See also Table S2 Figure S3.

To determine if human islet-derived glucagon was able to rescue phenotypes associated with glucagon loss in *NSG-GKO* host tissues, we examined *NSG-GKO Tx* mice for reversion of liver phenotypes. Fourteen weeks after human islet transplantation, we observed corrections of liver defects, including reduction of total and individual plasma amino acids (**Figures 3A-B and S3**), a reduction in liver glycogen levels (**Figure 3C**), and normalization of hepatic *G6pc, Tat, Oat*, and *Gls2* expression (**Figure 3D).** Together, these findings suggest that glucagon secretion by human islet grafts durably reconstituted a physiological islet-liver signaling axis.

### Glucagon secreted by human islet grafts corrects α cell hyperplasia in NSG-GKO mice

Impaired glucagon signaling in mice can evoke compensatory α cell proliferation and hyperplasia^19, 23, 24, 32^. Elevated circulating amino acids in mice lacking glucagon signaling were previously demonstrated to induce α cell proliferation and hyperplasia^20-22^ through a mechanism involving the α cell amino acid transporter *Slc38a5* ^20, 21^. As *NSG-GKO* mice exhibited hyperaminoacidemia, we next assessed islet α cell and β cell hyperplasia and proliferation in *NSG-GKO* islets. For α cell morphometry in *NSG-GKO* islets, we used a proglucagon-specific antibody that detected both wild type and internally-deleted GKO proglucagon (proglucagonΔ), and antibodies to detect MafB^33^, an adult α cell-specific islet transcription factor in mice (**Figures 4A-C**). Islet morphometry in adult *NSG-GKO* mice revealed an increased percentage of α cells expressing the proliferation marker Ki67 (**Figures 4E-H**) and increased α cell mass (**Figures 4A-D**). No difference was observed in β cell mass or proliferation in islets from *NSG-GKO* and *NSG* control mice (**Figures 4A-H**). In response to amino acids, mouse α cells proliferate and induce expression of Slc38a5^21, 22^, which is also expressed in mouse acinar cells^22, 34^. *NSG-GKO* mice showed an increase in α cell expression of Slc38a5 compared to islets from *NSG* control mice (**Figures 4I-N**), consistent with these prior studies. Thus, like in prior models of glucagon deficiency, we observed adaptive α cell expansion, stimulated by hyperaminoacidemia and accompanied by Slc38a5 expression.

**Figure 4.**
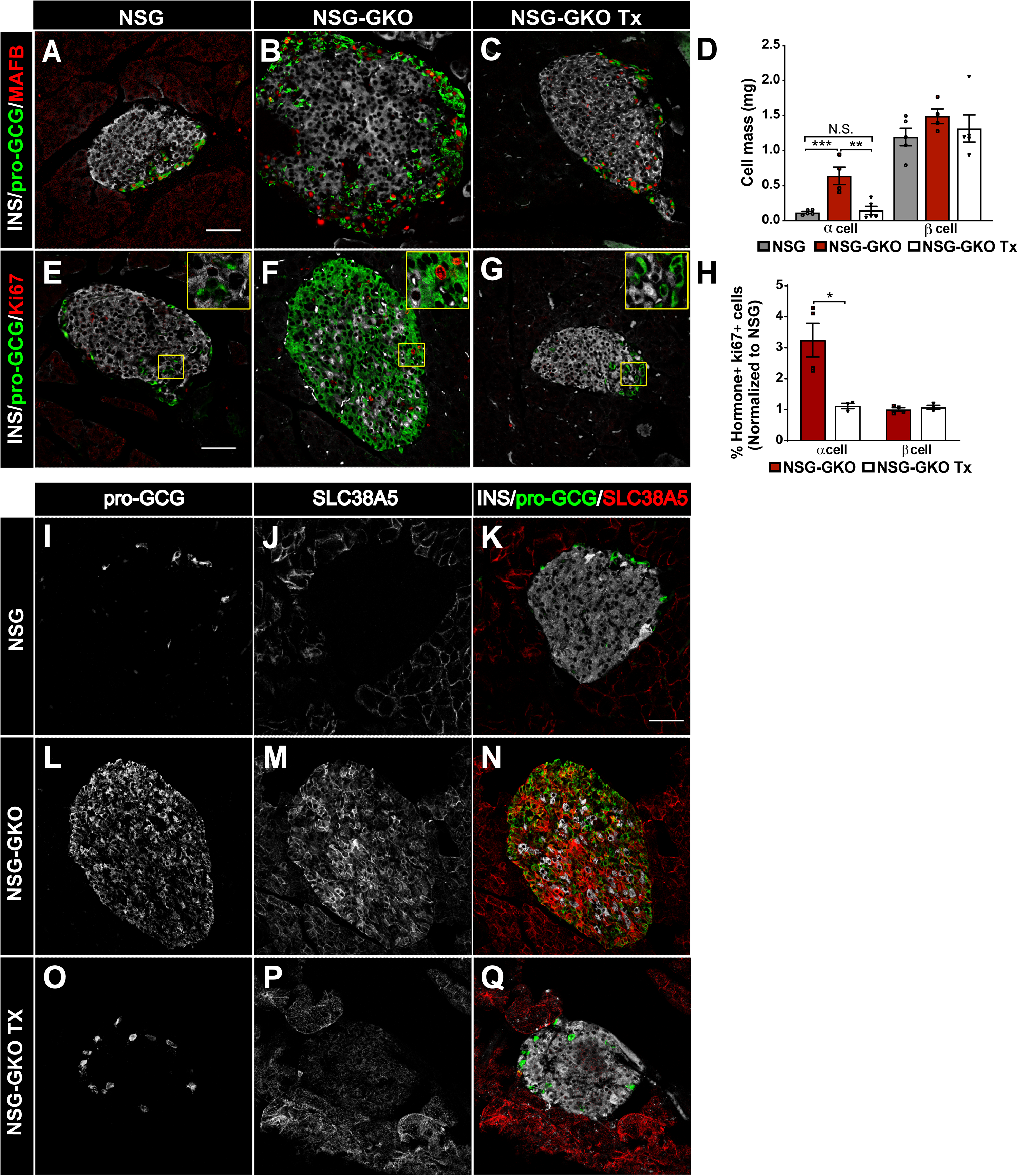
Human islet-derived glucagon corrects *NSG-GKO* α cell hyperplasia. (**A-C**) Representative immunostaining for morphometric analysis of *NSG* control, *NSG-GKO*, and *NSG-GKO* Tx mice with antibodies detecting proglucagon (green), Insulin (white), and MAFB (red). (**E-G**) Representative immunostaining for quantification of α and β cell proliferation in *NSG* control, *NSG-GKO*, and *NSG-GKO* Tx mouse pancreata using antibodies detecting proglucagon (green), insulin (white), and Ki67 (red). Quantification of α and β cell mass (**D**) (*NSG* vs. *NSG-GKO P*= 0.0010 and *NSG-GKO* vs. *NSG-GKO* Tx *P*= 0.0015 by one-way ANOVA, with Tukey’s multiple comparisons test) (*NSG* mice, n= 5; *NSG-GKO* mice, n= 4; *NSG-GKO* Tx mice, n= 5) and proliferation (**H**) (*P*= 0.0227 by two-tailed Student’s t-test) (*NSG* mice, n= 4; *NSG-GKO* mice, n= 4; *NSG-GKO* Tx mice, n= 3). (**I-Q**) Representative immunostaining of SLC38A5 expression in *NSG* control, *NSG-GKO*, and *NSG-GKO* Tx mouse pancreata using antibodies detecting proglucagon (green), insulin (white), and SLC38A5 (red). Scale bars, 50 **μ**m. Data are represented as mean of biological replicates ± SEM. * *P* ≤ 0.05, ** *P* ≤ 0.01, *** *P* ≤ 0.001.

As human glucagon from islet grafts restored circulating amino acid levels in *NSG-GKO* mice (**Figures 3A-B**), we next assessed the impact on host islet α cells in *NSG-GKO* Tx mice and controls. Morphometry revealed that host α cell mass ‘normalized’ in *NSG-GKO* Tx mice compared to *NSG* control mouse islets (**Figures 4C-D**), and was accompanied by a reduction in the number of proglucagonΔ^+^ Ki67^+^ cells (**Figures 4G-H**), and loss of Slc38a5 production in mouse α cells (**Figures 4O-Q**). Thus, restoration of glucagon signaling by human islet grafts in *NSG-GKO* mice corrected adaptive pancreatic islet α cell expansion observed in *NSG-GKO* mice.

### Restoration of glucose and insulin regulation in transplanted NSG-GKO mice

Glucagon increases blood glucose levels by promoting hepatic glucose output and is also implicated in regulating normal insulin secretion^35-37^. Hence, mice lacking glucagon signaling are hypoinsulinemic^18^ and hypoglycemic^19, 23, 38^. Consistent with these extant glucagon signaling mutant mouse models, *NSG-GKO* mice had chronically reduced blood glucose levels and lower *ad libitum* fed plasma insulin levels compared to *NSG* control mice (**Figures 5A-B and S4B-C**). Four weeks after human islet transplantation, *ad libitum* fed blood glucose and total plasma insulin levels in *NSG-GKO Tx* mice were increased and indistinguishable from *NSG* controls (**Figures 5A-B and S4B-C).** In *NSG-GKO Tx* mice, total plasma insulin levels reflected contributions from both host mouse β cells and transplanted human islets (**Figure S4D**). While circulating glucagon levels differed in *ad libitum* fed *NSG* and *NSG-GKO Tx* mice, human glucagon in *NSG-GKO Tx* mice was sufficient to maintain normoglycemia (**Figures 2B, 5A, and S4A-B**). Upon consideration of the appropriateness of glucagon secretion under acute hypoglycemia in *NSG-GKO Tx* mice (**Figure 2C**), these data suggest that human and mouse α cells may have distinct glycemic thresholds, supporting conclusions from other studies^39, 40^. Thus, transplanted human islets improved glycemic and insulin control in *NSG-GKO Tx* mice.

**Figure 5.**
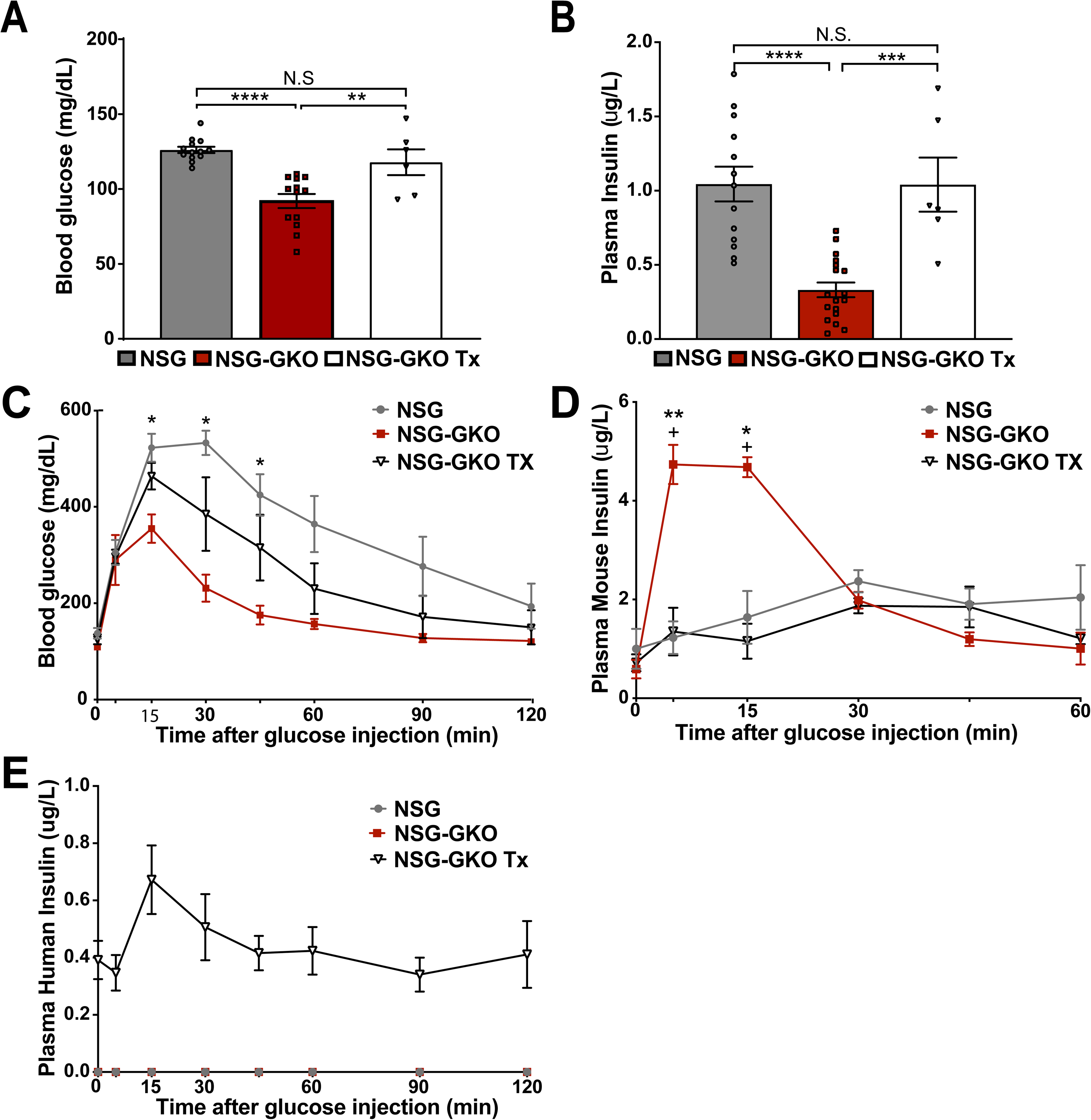
Improved glucose and insulin regulation in transplanted *NSG-GKO* mice. *NSG-GKO* mice post-transplantation (*NSG-GKO* Tx), *NSG-GKO* mice, and *NSG* control mice were assessed for ad libitum fed blood glucose (**A**) (*NSG* vs. *NSG-GKO P*≤ 0.0001 and *NSG-GKO* vs. *NSG-GKO* Tx *P*= 0.0038 by one-way ANOVA, with Tukey’s multiple comparisons test) (*NSG* mice, n= 13; *NSG-GKO* mice, n= 13; *NSG-GKO* Tx mice, n= 6) and plasma insulin levels (**B**) (*NSG* vs. *NSG-GKO P*≤ 0.0001 and *NSG-GKO* vs. *NSG-GKO* Tx *P*= 0.0003 by one-way ANOVA, with Tukey’s multiple comparisons test) (*NSG* mice, n= 13; *NSG-GKO* mice, n= 19; *NSG-GKO* Tx mice, n= 6). Mice were given an intraperitoneal glucose tolerance test and monitored for blood glucose measures (**C**) and plasma mouse (**D**) and human (**E**) insulin levels (*P* values generated by repeated measures ANOVA, with Tukey’s multiple comparisons test are listed in Table S3) (*NSG* mice, n= 3; *NSG-GKO* mice, n= 3; *NSG-GKO* Tx mice, n= 4). Human insulin excursion measured by human insulin-specific ELISA in the same IPGTT test as in panels C and D. Data are represented as mean of biological replicates ± SEM. * *NSG* vs. *NSG-GKO* mice, ^+^ *NSG-GKO* vs. *NSG-GKO* Tx mice (**C-D**). ^+^ or * *P* ≤ 0.05, ^++^ or ** *P* ≤ 0.01, ^+++^ or \*\*\**P* ≤ 0.001. See also Table S3 and Figure S4.

To examine insulin and glucose regulation further in *NSG-GKO* mice after human islet transplantation, we performed an intraperitoneal glucose challenge. Compared to *NSG* controls, glucose clearance by *NSG-GKO* mice was faster (**Figure 5C**) and accompanied by an exaggerated (mouse) insulin excursion (**Figure 5D**). Glucose and insulin excursions in *NSG-GKO* Tx mice more closely resembled that of *NSG* controls (**Figures 5C-D**). Dynamic total circulating insulin levels in *NSG-GKO* Tx mice reflected a combination of mouse and human insulins (**Figures 5D-E**). Notably, it appeared that human insulin release from transplanted islets was well-regulated during glucose challenge, leading to acute rise then clearance from the circulation (**Figure 5E**). These data suggest that human islet-derived glucagon improved glycemic and insulin regulation in *NSG-GKO* Tx mice, and thus highlight the role of glucagon in maintaining euglycemia and normal insulin secretion.

## Discussion

To address the absence of models to study regulated glucagon secretion from human islet α cells *in vivo*, here we used CRISPR/Cas9 to develop a glucagon-null immunocompromised mouse strain (*NSG-GKO*). Human islets engrafted durably in *NSG-GKO* mice and retained regulated glucagon and insulin secretion. Reconstituting glucagon signaling to target organs, like the pancreas and liver, rescued multiple phenotypes associated with glucagon deficiency, including deviations in circulating glucose, amino acid, and insulin levels. Prior reports of mice that lack glucagon signaling^18-23, 41, 42^ revealed many phenotypes associated with loss of glucagon signaling in pancreatic islets and other organs - phenotypes that we also observed in *NSG-GKO* mice. However, *NSG-GKO* mice also have distinct properties not previously reported^18, 23, 42^, likely reflecting preserved production of ‘nested’ proglucagon-derived peptides like GRPP, GLP-1, and GLP-2, in addition to the superior receptivity of the *NSG* strain background to xenotransplantation^6, 7^. While our approach led to the unavoidable loss of the oxyntomodulin and glicentin peptide hormones (which incorporate amino acids 1-29 of GCG), this also creates opportunities to study the *in vivo* functions of these hormones in *NSG-GKO* mice.

Glucagon is a crucial intercellular and inter-organ regulator of pancreatic islet cells, liver, and other organs^43^. Our results suggest that the *NSG-GKO* model should be useful for investigating these cell-signaling interactions. This proposal is based upon the observed reversal of hyperaminoacidemia, α cell hyperplasia, and islet Slc38a5 production in *NSG-GKO* mice after human islet transplantation, indicating (re)-establishment of at least two homeostatic *in vivo* signaling axes mediated by human glucagon and circulating amino acids. The first signaling axis links transplanted human islets to the host liver, and the second links the liver to native pancreatic islets cells. Further studies are needed to assess how other established intercellular inputs governing islet cell function and activity, like signaling from autonomic neurons, adrenal hormones, or afferents from the CNS^44^ might be reconstituted in the *NSG-GKO* model. Intra-islet signaling between α cells, β cells, δ cells, and other islet cells is known to regulate islet hormone secretion^45, 46^. Glucagon is a direct regulator of insulin secretion, while insulin inhibition of glucagon secretion is thought to be mediated through δ cell-derived somatostatin^35, 36, 47^. For example, we observed that restored circulating glucagon in *NSG-GKO* Tx mice curbed the excessive insulin secretion response to hyperglycemia seen in *NSG-GKO* mice, providing evidence that glucagon acts to restrain β cell post-prandial insulin secretion that may otherwise cause hypoglycemia. Thus, to the extent that regulated reciprocal interactions between human α cells and β cells are reconstituted and measurable in *NSG-GKO* mice, these mice could be useful for *in vivo* studies of these and other intra-islet signaling interactions. For instance, δ cell somatostatin is an inhibitor of glucagon secretion^47-49^ in rodents, and glucagon stimulates somatostatin secretion in perfused canine pancreas^50, 51^. However, human α and δ cell communication is largely uncharted. Studies of mechanisms governing cognate δ cell signaling in human islets should be feasible in the *NSG-GKO* model, a possibility requiring further investigation.

The *NSG-GKO* mouse allows powerful new ways to study human islets. To evaluate the function of candidate type 2 diabetes mellitus risk genes identified by GWAS and discover human islet β cell regulators, we previously used loss-, and gain-of-function genetics in human pseudo-islets transplanted in *NSG* mice^4, 10^. This experimental logic, using *NSG-GKO* mice, can now be expanded to human α cells in islets from previously-healthy, pre-diabetic, or diabetic donors. Importantly, we can now examine how α cell enriched genes^3, 52^ and genetic changes in diabetes mellitus^3, 53, 54^ impact human α cell identity and function. Moreover, we envision that *NSG-GKO* mice transplanted with islets from human donors (or other species) will be useful for examining how pharmacological agents, or acquired environmental stressors, like starvation or diet-induced obesity, impact human α cells. Using *NSG* mice, we recently reported that responses of transplanted human islet β cells to high fat diet challenge were distinct from those observed in (host) mouse β cells^9^. Additionally, *NSG-GKO* mice should be useful for assessing the function of transplanted islet-like cells produced from renewable sources like human stem cell lines^55-59^. Thus, *NSG-GKO* mice should be a useful resource for *in vivo s*tudies of human islets, islet replacement cells using genetics, small molecules, or modeling of acquired *in vivo* physiological or pathological risk states in diabetes mellitus.

## Acknowledgments

We thank past and current members of the Kim group for advice and encouragement, Dr. Sangbin Park for assistance in gene targeting, Dr. Owen McGuinness and the Vanderbilt University Medical Center Hormone Core (DK059637 and DK020593) for hormone measurements and advice, and Drs. David Serreze (JAX) and Kristin Abraham (NIDDK/NIH) for generation of mouse lines. This work was supported by the Type 1 Diabetes Mouse Resource (1UC4DK097610 to D. Serreze), a graduate research fellowship award from the National Science Foundation (DGF-114747 to K.T.), RO1 awards (DK107507; DK108817; CA21192701 to S.K.K.) and a UO1 award (DK120447 to Dr. P. MacDonald, Univ. of Alberta). Work in the Stein lab was supported by NIH grants (DK106755, DK050203, and DK090570 to R.W.S). Work in the Kim lab was also supported by NIH grant P30 DK116074, and by the Stanford Islet Research Core, and Diabetes Genomics and Analysis Core of the Stanford Diabetes Research Center.

## Author Contributions

Conceptualization: S.K.K., K.T. and Y.H.; Methodology: K.T. and Y.H., and S.K.K. Validation: K.T., Y.H., and X.G; Formal Analysis: K.T. and X.G.; Investigation: K.T., Y.H., and X.G.; Data Curation: K.T.; Writing - Original Draft: K.T. and S.K.K.; Writing - Review and Editing: K.T., Y.H., R.W.S., and S.K.K.; Visualization: K.T. and Y.H.; Supervision: R.W.S. and S.K.K; Project Administration: S.K.K.; Funding Acquisition: S.K.K.

## Methods

### Glucagon gene targeting in NSG mice

*NSG-GKO* mice were generated through the NIDDK Type 1 Diabetes Resource (TIDR) and the Jackson Laboratory. Two mouse *glucagon* Exon 3 guide RNAs (sgRNA 3569: GAAGACAAACGCCACTCACA and sgRNA 3572: CAGACTCTTACCGGTTCCTCT) and the CRISPR/Cas9 plasmid were injected into *NSG* oocytes to generate founders. Out of 33 progeny, one male (termed 18-1) was confirmed by TOPO cloning and DNA sequencing to carry the desired in-frame deletion and bred to *NSG* females for germ line transmission. Verified F1 heterozygous offspring were used for further breeding to homozygosity. Subsequent genotyping were performed using PCR amplification with 3573_F1 (TGAGGCTTTCCTAGTGCTGAG) and 3575_R1 (AACGATCAATACAGCTAAGGTCTC) primers, which produces a 715 bp wildtype product and 622 bp from the glucagon exon 3 deletion mutant (Figure 1B). Homozygous glucagon knockout mice were born at near Mendelian ratios. *NSG-GKO* mice were housed in a pathogen-free barrier facility at Stanford University Medical School, and were exposed to a normal 12-hour light cycle. Male and female mice (2-5 months old) were used for experiments along with age- and sex-matched control *NSG* mice.

### Human islet procurement and transplantation

Deidentified human pancreatic islets were procured through the Integrated Islet Distribution Program and Alberta Diabetes Institute Islet Core. Five hundred human islet equivalents (IEQ) from previously healthy, nondiabetic organ donors (n=6) with less than 15-hour cold ischemia time (Table 1) were used for transplantation under the kidney capsule of *NSG-GKO* mice as previously described^8,10^.

**Table 1.** Donor information of human islets used for transplantation studies.

| Sample ID | Age | Sex | BMI | Purity |
| --- | --- | --- | --- | --- |
| 1 | 40 | M | 25.5 | 95% |
| 2 | 57 | M | 27.6 | 80% |
| 3 | 31 | M | 31.8 | 80% |
| 4 | 47 | M | 27.6 | 90% |
| 5 | 60 | F | 30.6 | 80% |
| 6 | 30 | M | 25.5 | 90% |

### Glucose tolerance testing

After a 5-hour fast, mice were administered an intraperitoneal (IP) injection of D-glucose (3 g/kg body weight). Blood glucose levels were measured with a Contour glucometer (Bayer) at 0, 5, 15, 30, 45, 60, 90, and 120 minutes post injection and EDTA-treated plasma samples were collected for hormone assays. For circulating GLP-1 assessment, D-glucose (6 g/ kg body weight) was given by oral gavage and EDTA- and DPP4-inibitor-treated plasma samples were collected at 0, 15, and 30 minutes post-gavage.

### Insulin tolerance test (ITT)

After a 5-hour fast, mice were administered an intraperitoneal injection of Novolin R U-100 (1U/kg body weight). Blood glucose levels were measured with a Contour glucometer (Bayer) at 0, 15, and 30 minutes post insulin injection. EDTA- and protease cocktail-treated (Bimake) plasma samples were collected at 0 and 30 minutes post-insulin injection for circulating glucagon measurement.

### Plasma hormones and amino acids assays

Plasma insulin and glucagon levels were assessed using ultrasensitive mouse insulin ELISA (Mercodia) and glucagon ELISA (Mercodia), respectively. Circulating human insulin levels in transplanted *NSG-GKO* recipients were measured with human insulin ELISA (Mercodia). Plasma GLP-1 levels were quantified with an active GLP-1 ELISA (Eagle Biosciences). Plasma amino acid levels were determined using the L-Amino Acid Quantitation Kit (Sigma Aldrich) following manufacture’s instruction, and individual amino acids by the Vanderbilt University Hormone Assay and Analytical Services Core using a Biochrom 30 amino acid analyzer.

### Immunostaining and morphometry

Pancreata were weighed (wet weight), fixed in 4% paraformaldehyde overnight at 4°C, and 10 μm thick cryosections were prepared. At least 10 sections per pancreas spaced at least 100 μm apart were stained with the following antibodies: Guinea pig anti-insulin (Dako, 1:500), Rabbit anti-proglucagon (Cell Signaling Technologies, 1:400), Mouse anti-proglucagon (Novus Biologicals, 1:300), Guinea pig anti-glucagon (Takara, 1:2000), Rabbit anti-MafB (Bethyl, 1:250), Rat anti-Ki67 (Biolegend, 1:100), Rabbit anti-SLC38A5 (Abcam 1:200). Fluorescent micrographs were captured using a Zeiss AxioM1 microscope and a Leica SP2 confocal microscope. For islet mass analysis, fluorescent micrographs were captured with a 10X objective lens. For islet proliferation analysis and representative images presented in this manuscript, fluorescent micrographs were captured with a 40x objective lens. Images were processed in Image J for islet cell mass quantification and Image-Pro Plus for islet cell proliferation using previously described methods^60-61^.

### Liver glycogen content assessment

60 mg of tissue from the liver (left lateral lobe) was collected from mice fasted for 5 hours and flash frozen in liquid nitrogen. Liver samples were homogenized on ice in buffer containing protease cocktail inhibitor (Bimake) and quantified using a fluorometric glycogen assay kit according to manufacturer’s instructions (Cayman Chemicals).

### RNA isolation, cNDA synthesis, and quantitative PCR

90 mg of tissue from the liver (left lateral lobe) was collected from mice fasted for 5 hours. Then, total RNA was extracted using RNeasy Mini kit (Qiagen) and complementary DNA was synthesized using Maxima First Strand cDNA Synthesis kit (Thermo Fischer Scientific) following manufacturers’ instructions. Quantitative PCR (qPCR) was performed using TaqMan assays (Supplementary Table 1) and reagents from Applied Biosystems with *Actb* used as an endogenous control.

### Study approvals

All studies involving human islets were conducted in accordance with Stanford Unviersity Institutional Review Board guidelines. All animal experiments and methods were approved by and performed in accordance with the guidelines provided by Institutional Animal Care and Use Committee (IACUC) of Stanford University.

### Statistics

Data are presented as the mean of biological replicates ± SEM. All data are the result of one experiment per biological replicate, where each data point is a distinct biological replicate (n values listed in figure legends). GraphPad Prism v. 7 and Microsoft Excel were used to perform Student’s t-test (two-tailed), one-way ANOVA, repeated measures ANOVA, and two-way ANOVA statistical comparisons between data. *P* values ≤0.05 were considered significant.

## Data Availability

The data that support the findings of this study are available from the corresponding author upon request.

## Supplemental Information

### Figure Legends

**Figure S1.**
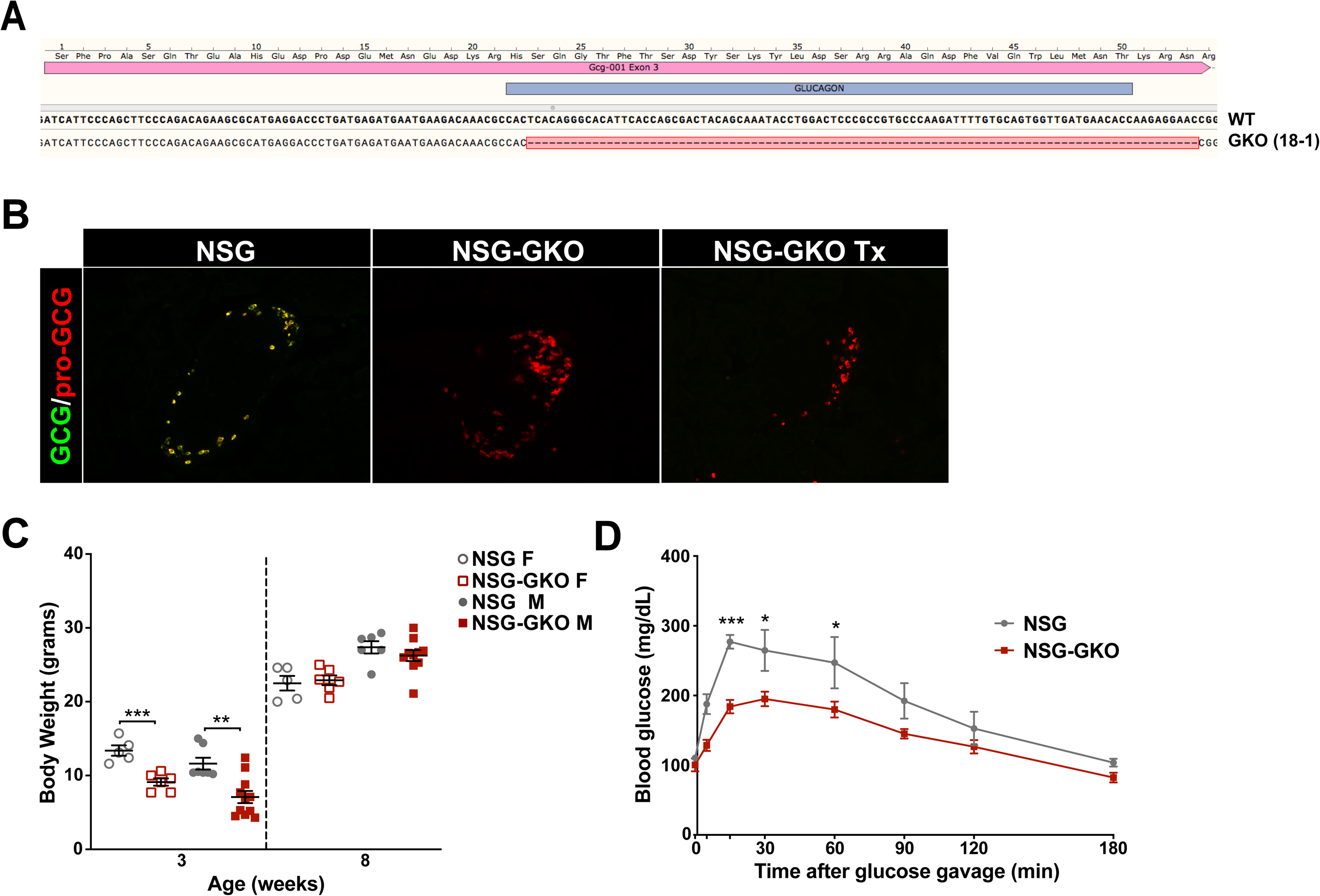
Related to Figure 1. (**A**) Sequence from *NSG-GKO* founder (18-1) showing an in-frame deletion of 93 base pairs within exon 3 of the *glucagon* gene compared to the WT-NSG sequence (WT). Pink bar on top depicts exon 3 of *glucagon*. Blue bar represents nucleotide sequences encoding mature glucagon peptide. Red-highlighted dashes indicate deleted nucleotides in founder 18-1. (**B**) Representative immunostaining of *NSG-GKO* pancreatic islets with antibodies raised against mature glucagon (GCG, green) and proglucagon (pro-GCG, red) - peptide sequences of GLP-1 (7-17). (**C**) Body weight of male and female *NSG-GKO* and *NSG* control littermates at 3 and 8-weeks of age (3-week old female mice P= 0.0007 and 3-week old male mice P= 0.0019 by two-tailed Student’s t-test) (*NSG* female mice, n= 5; *NSG-GKO* female mice, n= 6; *NSG* male mice, n= 7; *NSG-GKO* male mice, n= 11). (**D**) *NSG-GKO* and *NSG* control blood glucose measures over 180 minutes post oral glucose gavage (15’: P= 0.0007, 30’: P= 0.0185, 60’: P= 0.0259 by two-way ANOVA, with Tukey’s multiple comparisons test) (6g/kg body weight) (*NSG* mice, n= 4; *NSG-GKO* mice, n= 5). Data are represented as mean of biological replicates ± SEM. * *P* ≤ 0.05, ** *P* ≤ 0.01, *** *P* ≤ 0.001.

**Figure S2.**
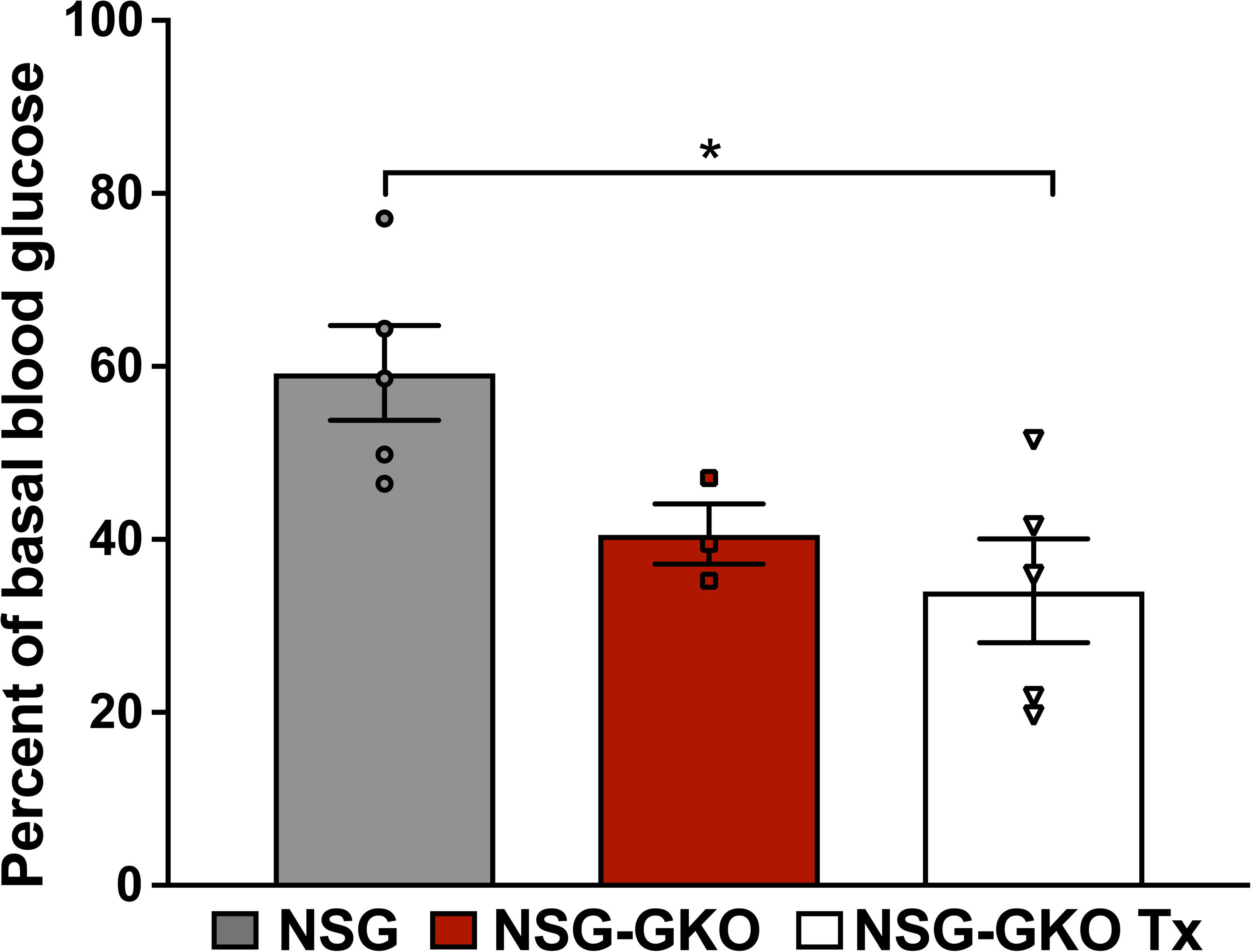
Related to Figure 2. Percent of basal blood glucose 30-minutes post insulin injection (1U/kg body weight) from *NSG, NSG-GKO, and NSG-GKO Tx* mice (P= 0.0017 by one-way ANOVA, with Tukey’s multiple comparison test) (*NSG* mice, n= 5; *NSG-GKO* mice, n= 3; *NSG-GKO* Tx mice, n= 5). Data are represented as mean of biological replicates ± SEM.

**Figure S3.**
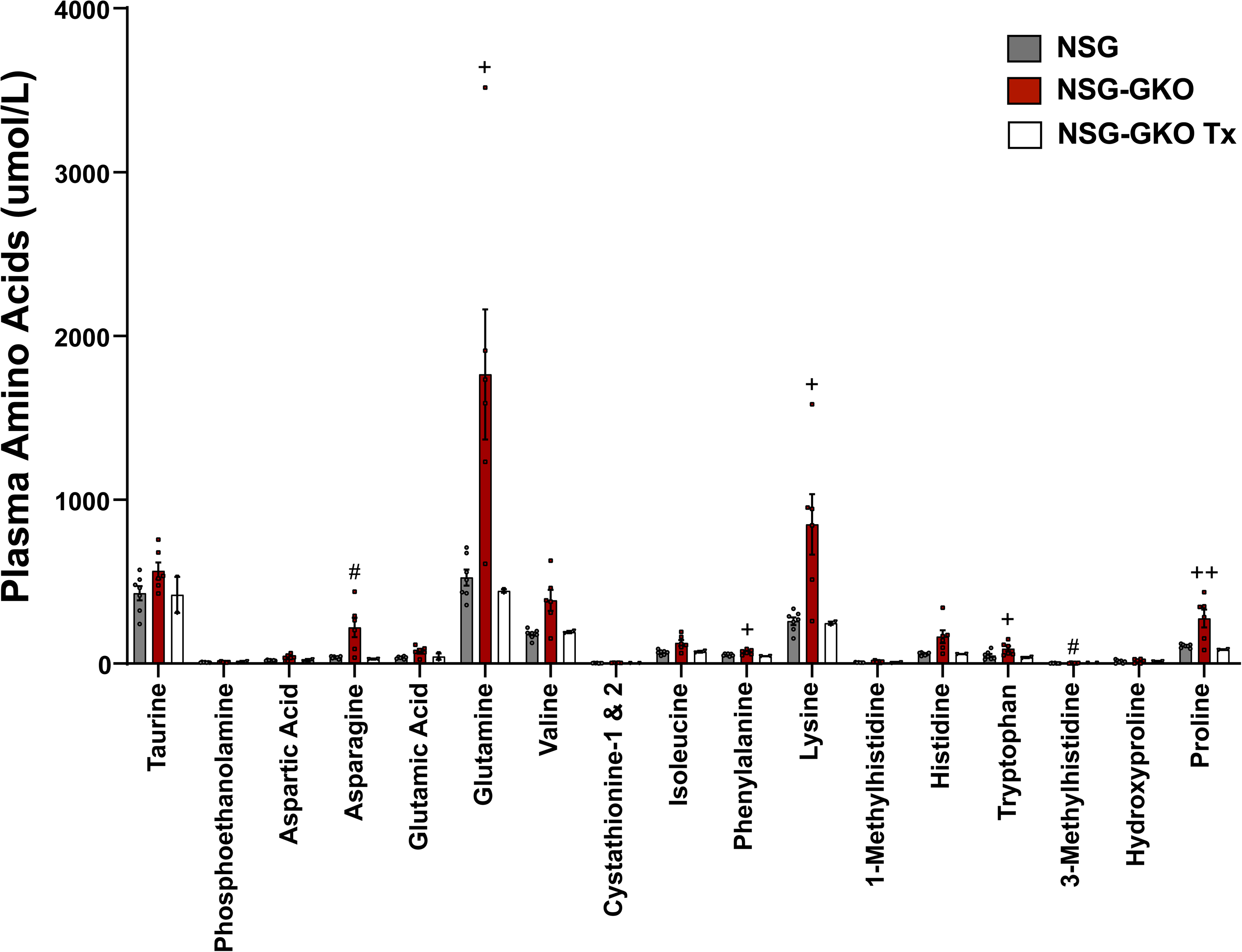
Related to Figure 3. Concentration of individual plasma amino acids that showed no significant changes in NSG-GKO mice (P values calculated by two-way ANOVA, with Tukey’s multiple comparison test are listed in Table S4) (*NSG* mice, n= 7; *NSG-GKO* mice, n= 6; *NSG-GKO* Tx mice, n= 2). Data are represented as mean of biological replicates ± SEM. ^+^ *NSG-GKO* vs. *NSG-GKO Tx* mice and ^#^ *NSG* vs. *NSG-GKO* Tx mice. ^#^ or ^+^ *P* ≤ 0.05, ^##^ or ^++^ *P* ≤ 0.01.

**Figure S4.**
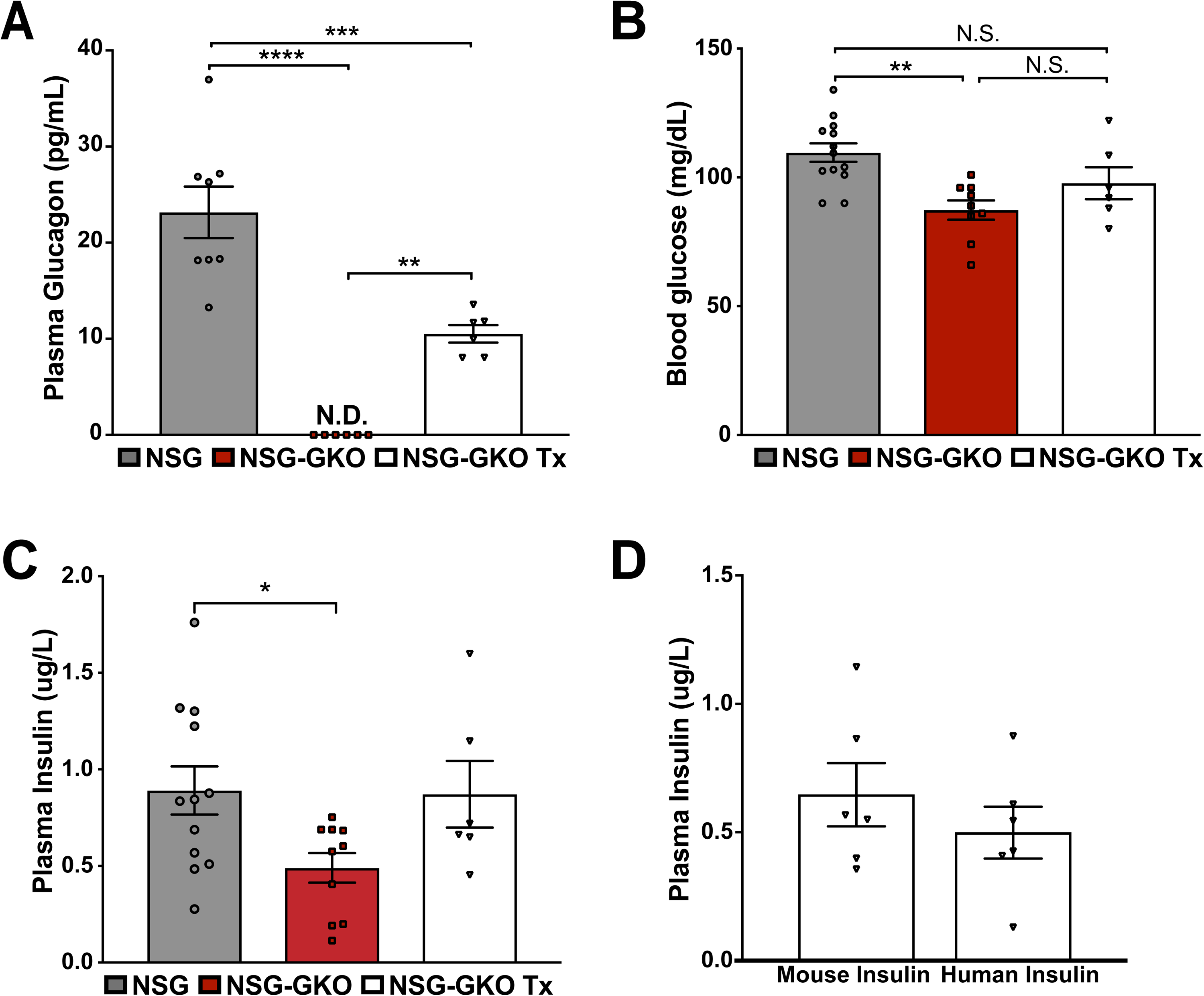
Related to Figure 5. Data are from *NSG* control, *NSG-GKO*, and *NSG-GKO* mice post-transplantation (*NSG-GKO* Tx). (**A**) Plasma glucagon levels in mice during *ad libitum* feeding (*NSG* vs. *NSG-GKO* P≤ 0.0001, *NSG-GKO* vs. *NSG-GKO* Tx P≤ 0.0023, and *NSG* vs. *NSG-GKO* Tx P= 0.004 by one-way ANOVA, with Tukey’s multiple comparison test) (*NSG* mice, n= 8; *NSG-GKO* mice, n= 6; *NSG-GKO* Tx mice, n= 7). Blood glucose (**B**) (P= 0.003 by one-way ANOVA, with Tukey’s multiple comparison test) (*NSG* mice, n= 13; *NSG-GKO* mice, n= 9; *NSG-GKO* Tx mice, n= 6) and plasma insulin levels (**C**) (P= 0.04 by one-way ANOVA, with Tukey’s multiple comparison test) (*NSG* mice, n= 12; *NSG-GKO* mice, n= 10; *NSG-GKO* Tx mice, n= 6) in fasted mice. (**D**) Mouse and human plasma insulin levels in ad libitum fed *NSG-GKO Tx* mice (n=6 mice). Data are represented as mean of biological replicates ± SEM. * *P* ≤ 0.05, ** *P* ≤ 0.01, *** *P* ≤ 0.001.

### Supplementary Tables

**Supplementary Table 1.**
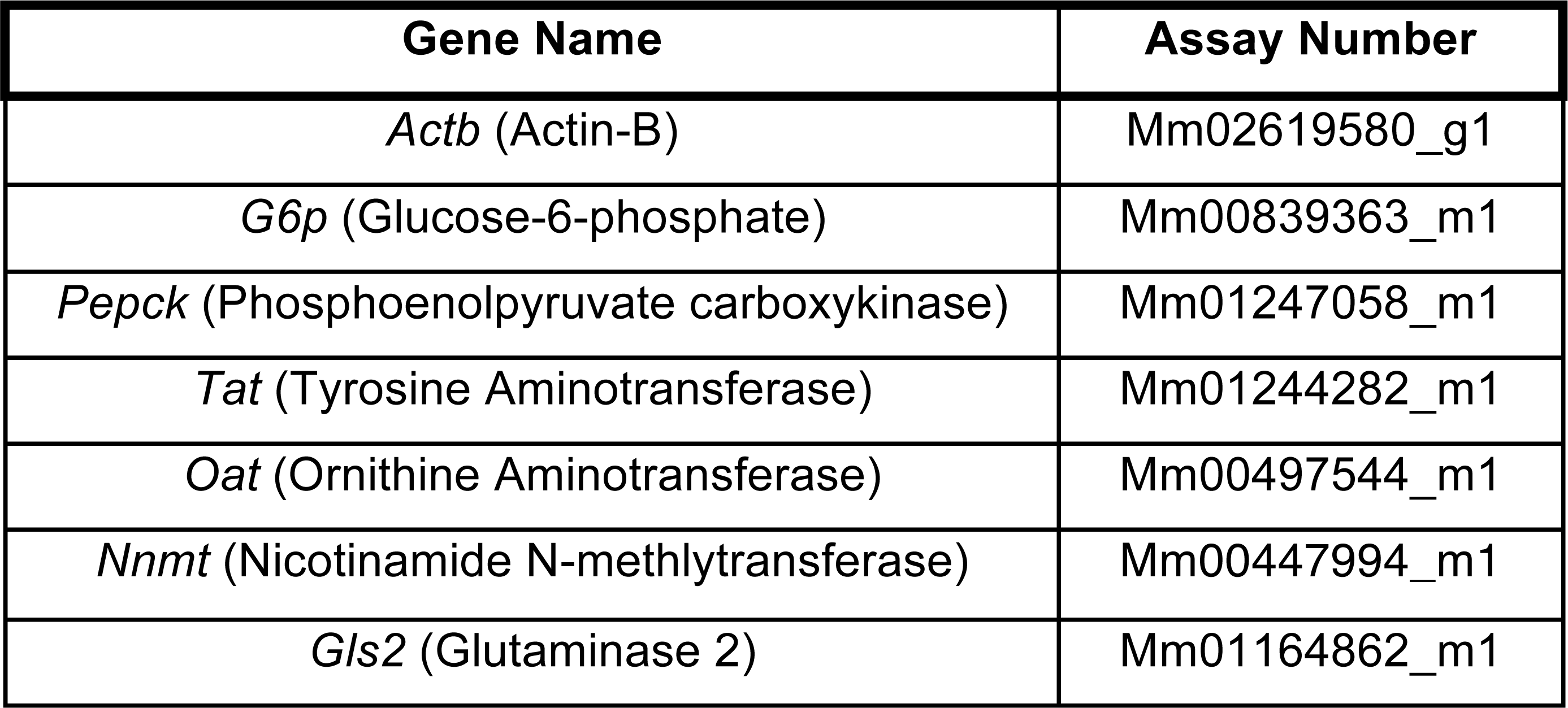
Taqman gene expression assay probes used in this study.

| Gene Name | Assay Number |
| --- | --- |
| *Actb* (Actin-B) | Mm02619580_g1 |
| *G6p* (Glucose-6-phosphate) | Mm00839363_m1 |
| *Pepck* (Phosphoenolpyruvate carboxykinase) | Mm01247058_m1 |
| *Tat* (Tyrosine Aminotransferase) | Mm01244282_m1 |
| *Oat* (Ornithine Aminotransferase) | Mm00497544_m1 |
| *Nnmt* (Nicotinamide N-methyltransferase) | Mm00447994_m1 |
| *Gls2* (Glutaminase 2) | Mm01164862_m1 |

**Supplementary Table 2.** P values for Figures 3B and 3D.

| Figure | Comparison | Adjusted P value |
| --- | --- | --- |
| 3B | **Threonine** |  |
|  | *NSG* vs. *NSG-GKO* | 0.0286 |
|  | *NSG-GKO* Tx vs. *NSG-GKO* | 0.0260 |
| 3B | **Serine** |  |
|  | *NSG* vs. *NSG-GKO* | 0.0442 |
|  | *NSG-GKO* Tx vs. *NSG-GKO* | 0.0420 |
| 3B | **Glycine** |  |
|  | *NSG* vs. *NSG-GKO* | 0.0422 |
|  | *NSG-GKO* Tx vs. *NSG-GKO* | 0.0334 |
| 3B | **Alanine** |  |
|  | *NSG* vs. *NSG-GKO* | 0.0470 |
|  | *NSG-GKO* Tx vs. *NSG-GKO* | 0.0405 |
| 3B | **Citrulline** |  |
|  | *NSG* vs. *NSG-GKO* | 0.0495 |
|  | *NSG-GKO* Tx vs. *NSG-GKO* | 0.0195 |
| 3B | **α-Amino-n-butyric Acid**<br>*NSG vs. NSG-GKO*<br>*NSG-GKO Tx vs. NSG-GKO* | 0.0362<br>0.0260 |
| 3B | **Cystine**<br>*NSG vs. NSG-GKO* | 0.0166 |
| 3B | **Methionine**<br>*NSG vs. NSG-GKO*<br>*NSG-GKO Tx vs. NSG-GKO* | 0.0294<br>0.0316 |
| 3B | **Leucine**<br>*NSG vs. NSG-GKO*<br>*NSG-GKO Tx vs. NSG-GKO* | 0.0450<br>0.0487 |
| 3B | **Tyrosine**<br>*NSG vs. NSG-GKO*<br>*NSG-GKO Tx vs. NSG-GKO* | 0.0315<br>0.0342 |
| 3B | **Ornithine**<br>*NSG vs. NSG-GKO*<br>*NSG-GKO Tx vs. NSG-GKO* | 0.0482<br>0.0393 |
| 3B | **Arginine**<br>*NSG vs. NSG-GKO*<br>*NSG-GKO Tx vs. NSG-GKO* | 0.0458<br>0.0405 |
| 3D | **G6pc**<br>*NSG vs. NSG-GKO* | 0.0237 |
| 3D | **Tat**<br>*NSG vs. NSG-GKO* | 0.0237 |
| 3D | **Gls2**<br>*NSG vs. NSG-GKO* | <0.0001 |
| 3D | **Oat**<br>*NSG vs. NSG-GKO* | 0.0003 |
| 3D | **Nmmt**<br>*NSG vs. NSG-GKO*<br>*NSG vs. NSG-GKO Tx* | <0.0001<br>0.0104 |

**Supplementary Table 3.** P values for Figures 5C and 5D.

| Figure | Comparison | Adjusted P value |
| --- | --- | --- |
| 5C | **15 minutes**<br>*NSG* vs. *NSG-GKO* | 0.0101 |
| 5C | **30 minutes**<br>*NSG* vs. *NSG-GKO* | 0.0133 |
| 5C | **45 minutes**<br>*NSG* vs. *NSG-GKO* | 0.0279 |
| 5D | **5 minutes**<br>*NSG* vs. *NSG-GKO*<br>*NSG-GKO* Tx vs. *NSG-GKO* | 0.0099<br>0.0231 |
| 5D | **15 minutes**<br>*NSG* vs. *NSG-GKO*<br>*NSG-GKO* Tx vs. *NSG-GKO* | 0.0116<br>0.0102 |

**Supplementary Table 4.** P values for Figure S4.

| Figure | Comparison | Adjusted P value |
| --- | --- | --- |
| S4 | **Asparagine**<br>*NSG* vs. *NSG-GKO* Tx | 0.0249 |
| S4 | **Glutamine**<br>*NSG-GKO* Tx vs. *NSG-GKO* | 0.0464 |
| S4 | **Phenylalanine**<br>*NSG-GKO* Tx vs. *NSG-GKO* | 0.0492 |
| S4 | **Tryptophan**<br>*NSG-GKO* Tx vs. *NSG-GKO* | 0.0462 |
| S4 | **3-Methylhistidine**<br>*NSG* vs. *NSG-GKO* Tx | 0.0174 |
| S4 | **Proline**<br>*NSG* vs. *NSG-GKO* Tx<br>*NSG-GKO* Tx vs. *NSG-GKO* | 0.0053<br>0.0380 |

